# ASD and ADHD have a similar burden of rare protein-truncating variants

**DOI:** 10.1101/277707

**Authors:** F. Kyle Satterstrom, Raymond K. Walters, Tarjinder Singh, Emilie M. Wigdor, Francesco Lescai, Ditte Demontis, Jack A. Kosmicki, Jakob Grove, Christine Stevens, Jonas Bybjerg-Grauholm, Marie Bækvad-Hansen, Duncan S. Palmer, Julian B. Maller, iPSYCH-Broad Consortium, Merete Nordentoft, Ole Mors, Elise B. Robinson, David M. Hougaard, Thomas M. Werge, Preben Bo Mortensen, Benjamin M. Neale, Anders D. Børglum, Mark J. Daly

## Main Text

### Introductory paragraph

Autism spectrum disorder (ASD) and attention-deficit/hyperactivity disorder (ADHD) are substantially heritable^1-4^, but individuals with psychiatric diagnoses often do not have blood drawn as part of routine medical procedure, making it difficult to collect large cohorts for genetic study. To overcome this challenge, we drew upon two Danish national resources: the Danish Neonatal Screening Biobank (DNSB) and the Danish national psychiatric registry. We have previously validated the use of archived bloodspots from the DNSB for genotyping^5^ and sequencing^6,7^, and we recently performed common variant analysis on dried bloodspot material in both ASD^8^ and ADHD^9^. Here, we present exome sequences from over 13,000 DNSB samples, finding that ASD and ADHD show a strikingly similar burden of rare protein-truncating variants, both significantly higher than controls. Additionally, the distributions of genes hit by these variants are not distinguishable between the two disorders, suggesting that many risk genes may be shared between them. These results motivate a combined analysis across ASD and ADHD, which—in conjunction with incorporation of the gnomAD reference database as additional population controls—leads to the identification of genes conferring general risk for childhood psychiatric disorders, including the novel gene MAP1A.

### Sample overview

Exome sequences for individuals included in the iPSYCH research initiative^10^ were obtained from archived dried blood samples stored by the DNSB. Individuals in this cohort were born in Denmark between 1981 and 2005. Using the Danish Psychiatric Central Research Register, we matched individuals to psychiatric diagnoses given by the end of 2012. After quality control, our dataset included sequences from 4,084 individuals with ASD, 3,536 individuals with ADHD, 727 individuals with both disorders, and 5,214 individuals designated as controls and without any of the primary psychiatric diagnoses studied in the iPSYCH initiative (ASD, ADHD, schizophrenia, bipolar disorder, affective disorder, anorexia) or any diagnosis of intellectual disability (ID) (Table 1). Although we allowed our cases to have more than one of the primary iPSYCH diagnoses (or ID), approximately 70% had no such documented comorbidity.

**Table 1:**
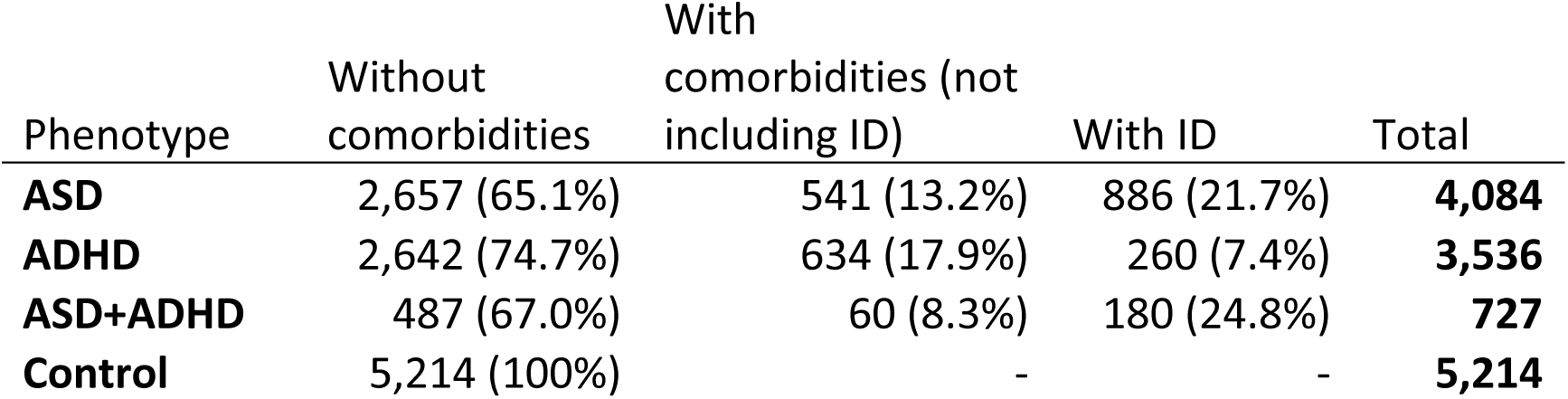
Phenotype breakdown of samples analyzed in this study. Percentages pertain to row totals. ID = intellectual disability.

### Rare variant rates

Studies of *de novo* variants in ASD have found that the greatest excess of point mutations carried by affected children resides in protein-truncating variants (PTVs; e.g., nonsense, frameshift, and essential splice site mutations)^11-14^. Furthermore, this excess burden is almost exclusively carried by PTVs that are not found in the Exome Aggregation Consortium (ExAC, http://exac.broadinstitute.org/) database and that occur in likely haploinsufficient genes (that is, genes with a probability of being loss-of-function intolerant, or pLI, of at least 0.9)^15,16^.

Without access to parental DNA in this study, we could not definitively determine whether a variant was a *de novo* mutation in the individual. We nonetheless followed studies of *de novo* variation in ASD as we attempted to define a set of filters to enrich for mutations that would be overrepresented in cases as compared to controls. Specifically, we tallied variants not found in ExAC and present in our dataset at 10 copies or fewer (which we define as “rare”) and that occurred in genes with a pLI of at least 0.9 (which we define as “constrained”). In samples without intellectual disability, we observed a significant excess of constrained rare PTVs (or “crPTVs”) in ASD cases (0.30/person, p=6.1E-14), ADHD cases (0.29/person, p=9.7E-11), and cases with both diagnoses (0.29/person, p=8.8E-04) compared to controls (0.21/person) (Figure 1). Consistent with previous observations, we also observed substantially higher rates of crPTVs in cases with comorbid ID compared to controls (0.41/person in ASD, p=6.7E-21; 0.35 in ADHD, p=8.4E-07; 0.43 in cases with both, p=5.0E-08) (Figure 1).

**Figure 1:**
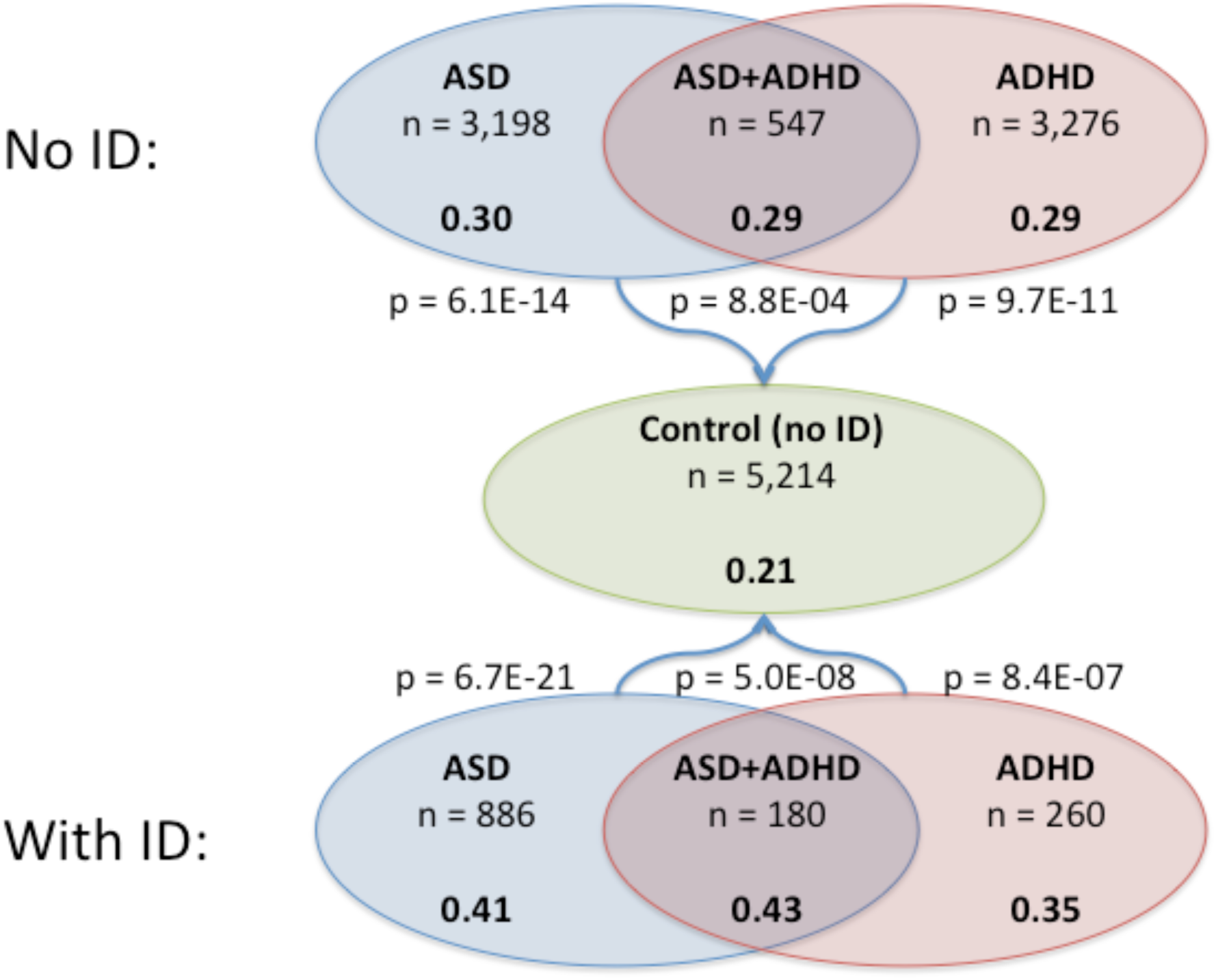
Per-person rates of constrained rare PTVs across phenotypes. P values are for comparison to controls by logistic regression.

In support of our filtering strategy, constrained PTVs found in ExAC (but present in our dataset at 10 copies or fewer) (Figure S1a), as well as rare PTVs in genes with pLI < 0.9 (Figure S1b), had p > 0.05 for all six case-control tests. Rates of constrained rare synonymous variation were similar across sample categories, both with and without ID (i.e., odds ratios compared to control ranged from 0.94 to 1.05), showing that the excess crPTVs in cases did not result from technical differences in variant calling (Figure S1c).

Across phenotypes, approximately 70%-75% of all crPTVs were found in people with exactly one crPTV, and the distributions within each case category were not significantly different from Poisson expectation (Figure S2), suggesting that the crPTVs occurred independently rather than being carried together in groups. Approximately 25-35% of cases had at least one crPTV (Table S1).

A similar trend to crPTVs was observed with rare missense variants, though the signal was less pronounced (e.g. 0.83/person in ASD cases without ID compared to 0.74 in Controls, p = 1.0E-04) (Figure S3). Here, we considered only missense variants with an MPC score (a measure of the deleteriousness of a missense variant based on a regional model of constraint^17^) of at least 2. Analogous to the crPTVs, little signal was observed when considering missense variants found in ExAC (but present in our dataset at 10 copies or fewer) (Figure S4a) or rare missense variants with MPC < 2 (Figure S4b).

### Comparison to published ASD data

To confirm the validity of the study and analysis design, we sought to compare the results of our case-control study with those previously seen in *de novo* studies of the Simons Simplex Collection (SSC) and Autism Sequencing Consortium (ASC) datasets^14,16^. Combining all of our cases with an ASD diagnosis (regardless of comorbidities), we observed a significantly enriched burden of rare PTVs in genes with three or more previously published *de novo* truncating variants in ASD (Table 2; p = 7.7E-03, OR = 3.0, n = 3,745 ASD cases without ID vs. 5,214 controls; p = 4.4E-11, OR = 16.6, n = 1,066 ASD cases with ID vs. 5,214 controls). The results become even more striking when disregarding *KDM5B*—the only gene in the set with *de novo* truncating variants observed in unaffected children—as this leaves 38 rare PTVs in our 4,811 cases and only a single rare PTV in 5,214 controls (Table 2; p = 9.2E-03, OR = 15.4, n = 3,745 ASD cases without ID vs. 5,214 controls; p = 1.9E-6, OR = 144.6, n = 1,066 ASD cases with ID vs. 5,214 controls). With such a strong signal in previously identified ASD genes, these results support the validity of Danish registry diagnoses and the quality of the bloodspots as a source of DNA.

**Table 2:** Rare PTV counts in genes with 3 or more published ASD *de novo* truncating variants. P values and odds ratios are for comparison to controls by logistic regression. OR = odds ratio. SE = standard error. Published data collected in references 14 and 16.

| Gene | Published *de novo* PTVs in ASD | Published *de novo* PTVs in unaffected children | Danish rare PTVs – ASD, no ID (n = 3,745) | Danish rare PTVs – ASD, ID (n = 1,066) | Danish rare PTVs – ASD, total (n = 4,811) | Danish rare PTVs – Control (n = 5,214) |
| --- | --- | --- | --- | --- | --- | --- |
| *CHD8* | 7 | 0 | 1 | 1 | 2 | 0 |
| *ARID1B* | 5 | 0 | 3 | 1 | 4 | 0 |
| *DYRK1A* | 5 | 0 | 0 | 2 | 2 | 0 |
| *SYNGAP1* | 5 | 0 | 0 | 4 | 4 | 0 |
| *ADNP* | 4 | 0 | 0 | 2 | 2 | 0 |
| *ANK2* | 4 | 0 | 5 | 3 | 8 | 1 |
| *DSCAM* | 4 | 0 | 1 | 0 | 1 | 0 |
| *SCN2A* | 4 | 0 | 1 | 3 | 4 | 0 |
| *ASH1L* | 3 | 0 | 0 | 2 | 2 | 0 |
| *CHD2* | 3 | 0 | 0 | 1 | 1 | 0 |
| *GRIN2B* | 3 | 0 | 0 | 4 | 4 | 0 |
| *POGZ* | 3 | 0 | 0 | 3 | 3 | 0 |
| *SUV420H1* | 3 | 0 | 1 | 0 | 1 | 0 |
| *KDM5B* | 3 | 2 | 7 | 1 | 8 | 8 |
| **Total, all genes** | 56 | 2 | 19 | 27 | 46 | 9 |
| OR vs Control | - | - | 3.0 | 16.6 | 5.9 | - |
| OR +/- SE | - | - | 2.0-4.6 | 10.8-25.4 | 4.0-8.5 | - |
| p | - | - | 7.70E-03 | 4.40E-11 | 1.70E-06 | - |
| **Total, no KDM5B** | 53 | 0 | 12 | 26 | 38 | 1 |
| OR vs Control | - | - | 15.4 | 144.6 | 41.6 | - |
| OR +/- SE | - | - | 5.4-44.0 | 50.9-410.2 | 15.1-114.6 | - |
| p | - | - | 9.20E-03 | 1.90E-06 | 2.30E-04 | - |

We also looked at Sanders et al. (2015)^14^ and asked if we had additional support for any of the genes with a putative association (e.g. 0.01 < FDR <= 0.1) with ASD in that study, and we found five genes for which we had at least two observed rare PTVs in ASD cases and none in controls (Table S2).

Further, the rate of crPTVs in the Danish data was similar to the combined rates of *de novo* and inherited crPTVs from the SSC+ASC data^16^ (Figure 2). The slightly higher rates in the Danish data are likely due in part to using newer exome capture and sequencing technology, enabling the calling of more variants overall. The similarity suggests that some of the signal captured by our case-control study is actually *de novo* variation, and that case-control studies can recover a class of variants that includes *de novo* mutations even when parents have not been sequenced.

**Figure 2:**
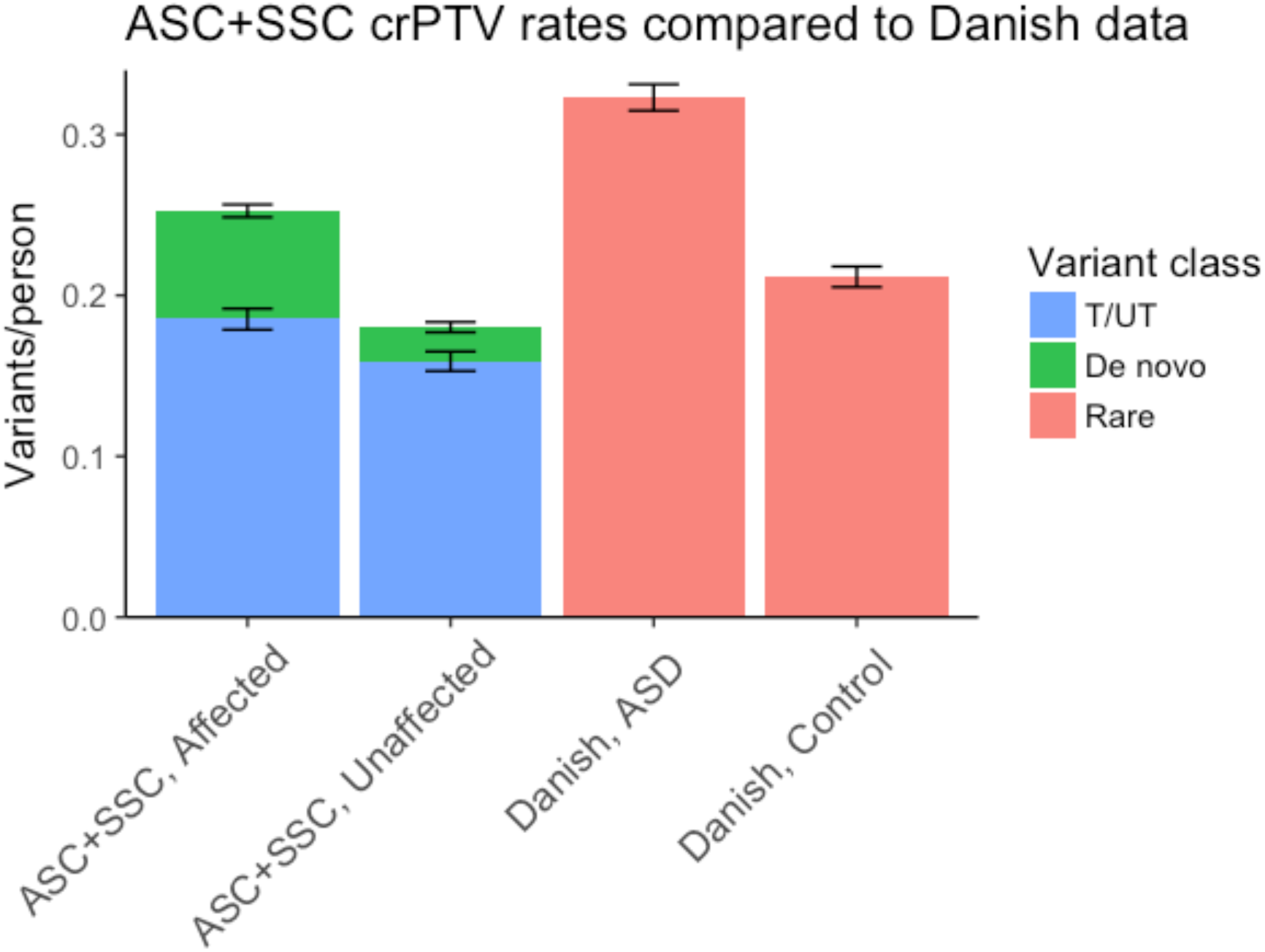
Comparing rates of constrained rare PTVs in Danish case-control data to SSC+ASC *de novo* and transmission data. SSC+ASC Affected is *de novo* variants in affected probands (n = 3,982) and transmitted variants (T) from parents of probands (n = 4,319). SSC+ASC Unaffected is *de novo* data from unaffected children (n = 2,078) and untransmitted (UT) variants from parents of probands (n = 4,319). Danish ASD data is crPTVs from all children with an ASD diagnosis (with or without ID, n = 4,811). SSC+ASC data from reference 16.

### Similarity of ASD and ADHD

Having observed similar rates of crPTVs between ASD and ADHD (without intellectual disability) (Figure 1), as well as several cases of ADHD with mutations in previously reported ASD genes (e.g., ADHD cases with rare PTVs in *ARID1B* and *ASH1L*), we decided to further explore the potential overlap of ASD and ADHD. We therefore focused on rates of rare PTVs occurring in the set of 455 genes with a published ASD *de novo* variant^11,12,16^. To rule out the possibility that a common comorbidity was driving the signal, we included only those cases with a single primary diagnosis and no intellectual disability (n = 2,657 for ASD and n = 2,642 for ADHD).

Strikingly, the ADHD cases had a slightly higher per-person rate than ASD cases of rare PTVs in the ASD-derived set of 455 genes (Figure 3). Both case categories were enriched above the control rate for rare PTVs (OR = 1.23 for ASD compared to control, p = 1.3E-02; OR = 1.24 for ADHD, p = 9.2E-03), but not for rare synonymous variants in the same genes (OR = 0.98 for ASD compared to control, p = 0.45; OR= for ADHD, p = 0.80). (Similar to the broader sample groups, the single-diagnosis ASD cases and ADHD cases had similar burdens of crPTVs overall, and both were significantly greater than controls, Figure S5.)

**Figure 3:**
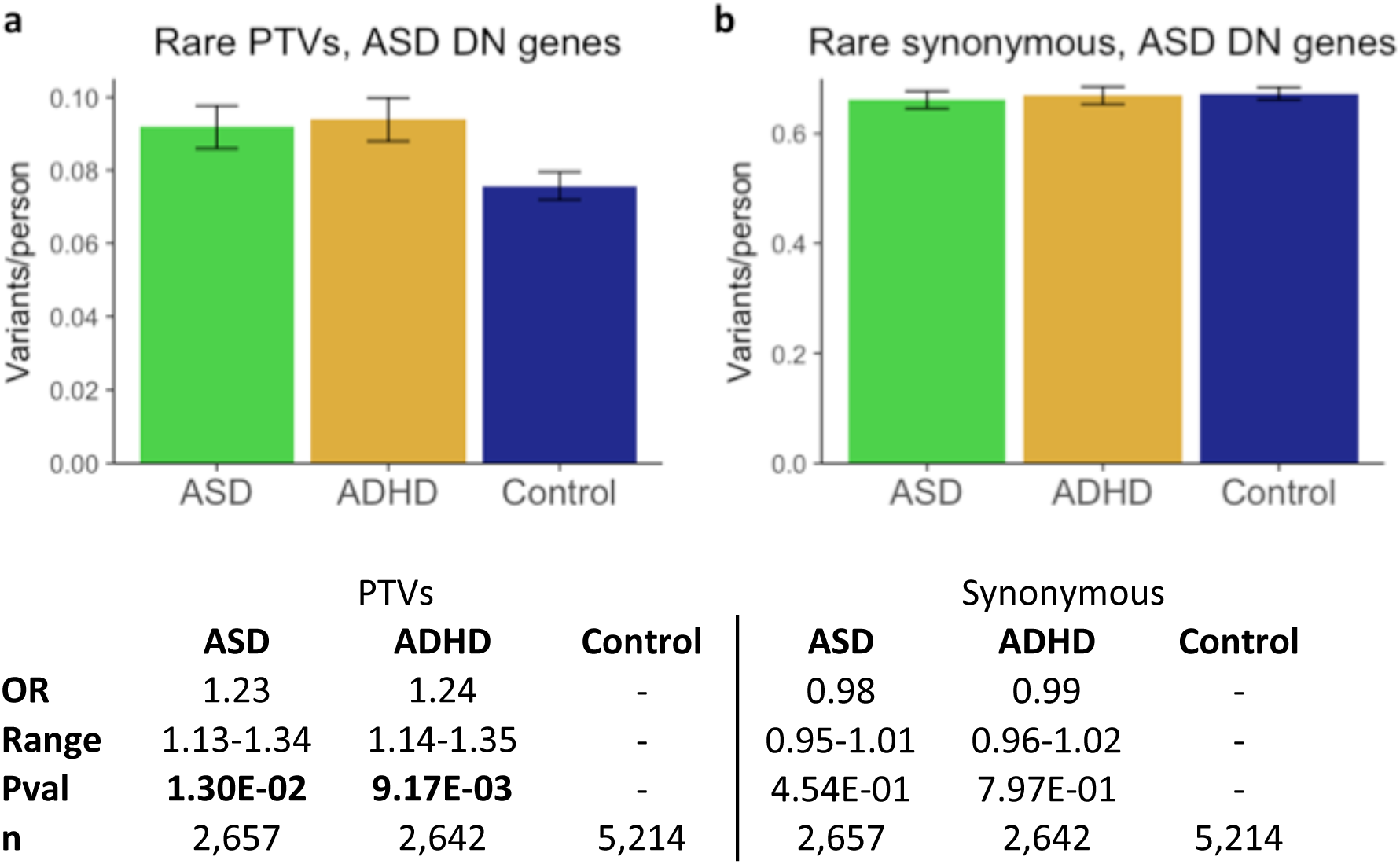
Rates of rare PTVs in 455 genes with a published ASD *de novo* mutation. a) Rare PTVs; b) Rare synonymous variants. ASD cases (n = 2,657) and ADHD cases (n = 2,642) were restricted to only those cases with a single primary diagnosis. P values are for comparison to controls (n = 5,214) by logistic regression. OR = odds ratio. Range = OR +/-standard error.

Given the similar rare PTV burden in ASD and ADHD cases, we used a c-alpha test to determine whether the genes with crPTVs were similar or distinct for ASD and ADHD. The c-alpha test^18^ can be used to test whether two distributions of rare variants have been selected from the same underlying distribution^19^. Considering again only cases with a single primary diagnosis, the test did not find a significant difference between ASD and ADHD, but it did find a significant difference when comparing either disorder and controls (Table 3). This result suggests that the crPTVs in people with ASD or ADHD are not only occurring at similar rates, but also in largely overlapping sets of genes. The test did not find a significant difference in any comparison when considering constrained rare synonymous variation.

**Table 3:** c-alpha test results.

| Comparison | PTVs |  | Synonymous variants |  |
| --- | --- | --- | --- | --- |
|  | Genes<br>(>1 event) | p value | Genes<br>(>1 event) | p value |
| **ASD vs ADHD** | 333 | 0.83 | 2865 | 0.086 |
| **ASD vs Control** | 438 | 4.4E-07 | 2987 | 0.11 |
| **ADHD vs Control** | 428 | 2.4E-05 | 2982 | 0.88 |

### Most-hit genes when combining case categories

The unexpected finding that ASD and ADHD had similar burdens of crPTVs occurring in similar genes motivated pooling ASD and ADHD cases (n = 8,347) for the purposes of gene discovery. To increase our control population, we included non-Finnish Europeans from the non-psychiatric subset of gnomAD (n = 43,836) (http://gnomad.broadinstitute.org), for a total of 49,050 controls.

To ensure that these cohorts were comparable, we determined the regions of the exome that were well-covered (80% of the samples had at least 10x coverage) in both the Danish exomes and the gnomAD exomes, and we only considered variants in these regions. We ruled out all variants present at more than 5 copies in the combined 57,397-person dataset, and we then counted the number of protein-truncating and synonymous variants by gene. We used Fisher’s exact test to calculate case vs control p values for PTV and synonymous counts in each gene, and we focused on genes with equal or greater truncating variation in cases than controls. To ensure that the comparison was not biased by differential sequencing quality between cases and controls, we excluded all genes in which synonymous variation rates were higher in cases than controls at a nominal threshold of p < 0.05. The vast majority of genes in fact had higher rates of synonymous variation in controls than in cases, indicating that more variants were, on average, being called per sample in gnomAD (potentially due to factors including DNA source and QC methodology)—a trend which any PTV had to overcome in order to have a greater burden in cases than controls.

The top result in this analysis was microtubule-associated protein 1A (*MAP1A*), in which we observed 12 PTVs in Danish cases (4 ASD without ID, 3 ASD with ID, 5 ADHD without ID), none in controls, and only 4 in gnomAD (p = 9.1E-08) (Table 4; Supplementary Data). *MAP1A* is highly expressed in the mammalian brain and is important for organization of neuronal microtubules; a candidate gene study identified an excess of rare missense variants in *MAP1A* in ASD and schizophrenia^20^. Our second result, *DYNC1H1*, had 8 case PTVs (4 ASD without ID, 2 ASD with ID, 1 ASD+ADHD with ID, 1 ADHD without ID) and has also been implicated in neurological disorders^21^. Among genes with a p value of less than 0.01, we observed three genes previously associated with ASD by Sanders et al. (2015)^14^ – *POGZ, SCN2A*, and *ANK2*. *ANK2* had 7 case PTVs (1 ASD without ID, 2 ASD with ID, 2 ASD+ADHD without ID, 1 ASD+ADHD with ID) and is particularly noteworthy, as this gene has a higher burden of published *de novo* mutations in ASD than it does in intellectual disability. We also identified an excess of rare PTVs in several known ID/DD genes (e.g., *ANKRD11, CTNNB1, MEF2C*), and indeed cases with mutations in these genes had high rates of ID (e.g., all four individuals with a PTV in MEF2C also have a diagnosis of ID), suggesting that diagnoses of ASD or ADHD were components of a broader neurodevelopmental disorder in these instances. A quantile-quantile plot is shown in Figure S6a, and an analogous plot for synonymous variants (Figure S6b) shows little inflation.

**Table 4:** Top genes by constrained rare PTV count, ranked by Fisher’s exact p value. (Note that *SCN2A* has 4 PTVs listed in Table 2 but only 3 listed here because one fell 2bp outside the consensus high-confidence region used when combining the Danish data with gnomAD.) Showing only genes with pLI ≥ 0.9.

| Gene | Case<br>(n = 8,347) | Control<br>(n = 5,214) | p value<br>(Danish) | gnomAD<br>(n = 43,836) | p value<br>(Comb.) | OR | ASD<br>*dn* | DDD<br>*dn* | Prev.<br>FDR |
| --- | --- | --- | --- | --- | --- | --- | --- | --- | --- |
| *MAP1A* | 12 | 0 | 4.92E-03 | 4 | 9.10E-08 | 17.7 | 0 | 1 | - |
| *DYNC1H1* | 8 | 0 | 2.72E-02 | 5 | 1.29E-04 | 9.4 | 0 | 0 | - |
| *ANKRD11* | 5 | 0 | 1.64E-01 | 1 | 3.43E-04 | 29.4 | 2 | 32 | - |
| *ZNF536* | 4 | 0 | 3.05E-01 | 0 | 4.47E-04 | Inf | 0 | 0 | - |
| *SPTBN1* | 5 | 0 | 1.64E-01 | 2 | 1.05E-03 | 14.7 | 1 | 1 | - |
| *TM9SF4* | 5 | 0 | 1.64E-01 | 2 | 1.05E-03 | 14.7 | 0 | 0 | - |
| *TNRC6C* | 7 | 0 | 4.86E-02 | 7 | 1.83E-03 | 5.9 | 0 | 0 | - |
| *POGZ* | 4 | 0 | 3.05E-01 | 1 | 1.98E-03 | 23.5 | 3 | 12 | < 0.01 |
| *RAI1* | 5 | 0 | 1.64E-01 | 3 | 2.47E-03 | 9.8 | 1 | 2 | - |
| *CTNNB1* | 3 | 0 | 2.90E-01 | 0 | 3.07E-03 | Inf | 1 | 15 | - |
| *GLUL* | 3 | 0 | 2.90E-01 | 0 | 3.07E-03 | Inf | 0 | 0 | - |
| *HCN1* | 3 | 0 | 2.90E-01 | 0 | 3.07E-03 | Inf | 0 | 0 | - |
| *SCN2A* | 3 | 0 | 2.90E-01 | 0 | 3.07E-03 | Inf | 4 | 10 | < 0.01 |
| *STAT5B* | 3 | 0 | 2.90E-01 | 0 | 3.07E-03 | Inf | 0 | 0 | - |
| *ZNF445* | 3 | 0 | 2.90E-01 | 0 | 3.07E-03 | Inf | 0 | 0 | - |
| *HSPA12A* | 6 | 1 | 2.62E-01 | 5 | 3.95E-03 | 5.9 | 0 | 0 | - |
| *KIAA1549* | 8 | 1 | 1.67E-01 | 11 | 4.87E-03 | 3.9 | 0 | 0 | - |
| *GRM3* | 4 | 0 | 3.05E-01 | 2 | 5.24E-03 | 11.8 | 0 | 0 | - |
| *TSC22D1* | 4 | 0 | 3.05E-01 | 2 | 5.24E-03 | 11.8 | 0 | 0 | - |
| *ZNF407* | 4 | 0 | 3.05E-01 | 2 | 5.24E-03 | 11.8 | 0 | 0 | - |
| *LRRC25* | 4 | 1 | 6.55E-01 | 1 | 5.24E-03 | 11.8 | 0 | 0 | - |
| *ANK2* | 7 | 1 | 1.64E-01 | 9 | 6.96E-03 | 4.1 | 4 | 1 | < 0.01 |
| *GPR116* | 9 | 2 | 2.22E-01 | 15 | 8.65E-03 | 3.1 | 0 | 0 | - |
| *MYCBP2* | 6 | 2 | 7.19E-01 | 6 | 9.91E-03 | 4.4 | 0 | 0 | - |
| *NR3C2* | 4 | 0 | 3.05E-01 | 3 | 1.08E-02 | 7.8 | 1 | 1 | - |

We conducted a similar analysis of missense variants with an MPC score of at least 2 (Table S3; Figure S6c; Supplementary Data). In this analysis, our top hit was *PHOX2B* (p = 5.8E-05), a gene involved in the formation of the autonomic nervous system during development^22^.

## Discussion

In summary, we have shown that archived bloodspots can be used to generate good quality exome sequencing data and that the risk for ASD driven by crPTVs in the Danish exome data is broadly consistent with that previously seen in SSC+ASC data. Perhaps surprisingly, we observed a similar burden of rare PTVs in ASD and ADHD, both in constrained genes overall and in genes with a previously published ASD *de novo* mutation, and we further showed a clear overlap in the rare variant signal between the two disorders.

With careful matching of sequence depth and variation rates, we used gnomAD as an additional control population and identified *MAP1A* as significantly associated with ASD and ADHD. *MAP1A* is a gene that has not been flagged by previous studies of intellectual disability or developmental delay, and it may represent genes where truncating variants are relevant to psychiatric cases (across diagnostic boundaries) with milder or more behavioral profiles in addition to those characterized by more profound neurodevelopmental symptomatology.

Genetic connections between ASD and ADHD have been made previously; for example, twin studies show that traits related to ASD significantly co-occur with traits related to ADHD^23,24^, suggesting a shared genetic component to their etiology. Siblings of children with an ASD diagnosis are also more likely to exhibit symptoms of ADHD and develop ADHD than the general population^25,26^. In addition, rare copy number variants have been found in some of the same genes in individuals with ADHD as in individuals with ASD^27^, and CNVs in autism and ADHD occur in the same biological pathways more often than would be expected by chance^28^. In the genotype data from our population sample, additional compelling evidence comes from the highly non-random rate of comorbidity and the recent finding of shared polygenic risk based on common variants^8^. This study, however, is the first to have such a large sample size of exome sequences to analyze in the two disorders, enabling comparisons such as the c-alpha test.

Of importance moving forward, the similar burden of crPTVs in ASD and ADHD suggests that study designs that have been successful in ASD will also be useful in ADHD. We also provide strong support for the idea that combining studies across childhood disorders may increase power and provide a more complete picture of the phenotypic consequence of rare variation in each gene.

## Online Methods

### How samples were selected

Individuals in the iPSYCH cohort were born in Denmark between May 1, 1981 and December 31, 2005. Neonatal dried blood samples were stored in the Danish Neonatal Screening Biobank. The iPSYCH initiative considers six primary psychiatric diagnoses—autism, ADHD, schizophrenia, bipolar disorder, affective disorder, and anorexia—and individuals were selected for inclusion in the cohort after matching them to psychiatric diagnoses in the Danish Psychiatric Central Research Register. Autism cases include individuals with an ICD10 diagnosis code of F84.0, F84.1, F84.5, F84.8, or F84.9. ADHD cases include individuals with an F90.0 diagnosis. The intellectual disability designation was based on an individual having any diagnosis for intellectual disability, including mild, moderate, or severe (codes F70-F79). The study was approved by the Regional Scientific Ethics Committee in Denmark and the Danish Data Protection Agency.

### How samples were sequenced

The primary iPSYCH cohort consists of over 88,000 samples. After DNA extraction by the DNSB and sample genotyping, a subset of approximately 20,000 age- and ancestry-matched samples was selected for exome sequencing. These samples had phenotypes across the range of disorders studied in the iPSYCH research initiative. Sequencing was performed by the Genomics Platform of the Broad Institute in Cambridge, MA, using an Illumina Nextera capture kit and an Illumina HiSeq sequencer. Sequencing was carried out in multiple waves, including a smaller pilot wave (“Pilot 1”) and two larger production waves (“Wave 1” and “Wave 2”).

### How sequencing data was processed to create callset

Raw sequencing data was processed using the Genome Analysis Toolkit^29^ (GATK) version 3.4 to produce a VCF version 4.1 variant callset file. The VCF used as the starting point for this study included 586 samples from Pilot 1, 6,733 samples from Wave 1, and 12,532 samples from Wave 2.

### How callset was QC’d

Most filtering steps downstream of GATK were performed in the scalable genomics program Hail (https://hail.is). Principal component analysis on a set of 5,848 common variants was used to restrict the analysis to only putatively European samples (removed 1,924/19,851 samples). Duplicate samples (5 samples) and samples where comparison of reported sex to imputed sex indicated potential Turner (1 sample) or Klinefelter (8 samples) syndromes were removed. Samples with levels of chimeric reads or estimated contamination above 5% (61 samples) were then removed from the dataset. Variants were removed if they did not pass GATK variant quality score recalibration or if they fell outside the exome target.

Several genotype filters were used to remove calls of low quality:

- Any call with a depth a) less than 10 or b) greater than 1000;
- Homozygous reference calls with a) GQ less than 25 or b) less than 90% reads supporting the reference allele;
- Homozygous variant calls with a) PL(HomRef) less than 25 or b) less than 90% reads supporting the alternate allele;
- Heterozygote calls with a) PL(HomRef) less than 25, b) less than 25% reads supporting the alternate allele, c) fewer than 90% informative reads (e.g. number of reads supporting the reference allele plus number of reads supporting the alternate allele less than 90% of the read depth), d) a probability of drawing the allele balance from a binomial distribution centered on 0.5 of less than 1e-9, or e) a location where the sample should be hemizygous (e.g. calls on the X chromosome outside the pseudoautosomal region in a male).
- Any call on the Y chromosome outside the pseudoautosomal region on a sample from a female.

Following the application of these genotype filters, three call rate filters were used: first the removal of variants with a call rate below 90%, then the removal of samples with a call rate below 95% (580 samples), then the removal of variants with a call rate below 95%. Between the sample call rate filter and the final variant call rate filter, one of each pair of related samples was removed (124 samples). Relatedness was calculated using the program KING^30^, and relatedness was defined as a relationship value of 0.1 or greater (approximately corresponding to a pi-hat of 0.2). Variants remaining in the dataset were annotated with the Variant Effect Predictor^31^, and one transcript for each variant was selected (prioritizing canonical coding transcripts) to assign a gene and a consequence to each variant.

Following the application of these filters, the dataset contained 17,148 individuals, some of whom had phenotypes not analyzed in the work presented here. The MAC value used when applying the 10-copy threshold for “rare” variant analyses was calculated within this group.

As a final quality control step, samples were removed (169 samples) if they fell in the top 1% of samples by number of not-in-ExAC singletons, as this added noise was an indicator that their quality may have decayed with the age of the archived sample. The remaining ASD (4,084), ADHD (3,536), ASD+ADHD (727), and control (5,214) samples were the ones used in this study.

All references to ExAC in this study refer to release 0.3 of the non-psychiatric (“nonpsych”) subset which has had samples from psychiatric studies removed (ftp://ftp.broadinstitute.org/pub/ExAC_release/release0.3/subsets/ExAC.r0.3.nonpsych.sites.vcf.gz).

### P value calculations

For calculating p values for classes of variants (e.g. crPTVs), covariates included in the logistic regression model were birth year, sex, the first ten principal components (of PCA carried out after dropping non-European samples), number of synonymous variants passing frequency filters (both in relevant pLI category and total), percent of exome target covered at a read depth of at least 20, mean read depth at sites within exome target passing VQSR, number of SNPs (of any population frequency) at sites within exome target passing VQSR, library capture set, and sequencing wave. For calculating gene-level p values, Fisher’s exact test was used. For both calculations, PTV counts were capped at one per person per gene to correct for the rare situation where one insertion or deletion event is labeled as two separate frameshifts by the genotype caller.

For Figure S2, the R function chisq.test was used with observed frequencies and Poisson-expected probabilities based on the observed mean. P values were simulated with 10,000 replicates.

### c-alpha

The c-alpha tests utilized the R package AssotesteR (http://cran.r-project.org/web/packages/AssotesteR/index.html), with PTVs again capped at one per person per gene. These tests did not consider genes with only one PTV across individuals. We ran 10,000 permutations and checked that the permutation-based p value was not larger than the reported analytic p value.

### q-q plots

The PTV q-q plot (Figure S6a) included all genes which 1) had at least two PTVs across Danish case and control samples, 2) did not have an odds ratio greater than 1 for synonymous variants (with corresponding p value smaller than 0.1) when comparing cases to the combined Danish and gnomAD controls, and 3) had an odds ratio greater than 1 for PTVs when comparing cases to the combined Danish and gnomAD controls. Of the 5,506 genes that passed the first filter, only 29 failed the second. A total of 2,370 genes passed all three filters. The missense q-q plot (Figure S6c) was generated analogously.

The synonymous q-q plot (Figure S6b) included all genes which 1) had at least two synonymous variants across Danish case and control samples and 2) had an odds ratio greater than 1 for synonymous variants when comparing cases to the combined Danish and gnomAD controls. Of the 15,318 variants that passed the first filter, only 1,679 also passed the second.

## Acknowledgements

The iPSYCH project is funded by the Lundbeck Foundation (grant numbers R102-A9118 and R155-2014-1724) and the universities and university hospitals of Aarhus and Copenhagen. The Danish National Biobank resource at the Statens Serum Institut was supported by the Novo Nordisk Foundation. Sequencing of iPSYCH samples was supported by grants from the Simons Foundation (SFARI 311789 to MJD) and the Stanley Foundation. Other support for this study was received from the NIMH (5U01MH094432-02, 5U01MH111660-02, and U01MH100229 to MJD). Computational resources for handling and statistical analysis of iPSYCH data on the GenomeDK and Computerome HPC facilities were provided by, respectively, Centre for Integrative Sequencing, iSEQ, Aarhus University, Denmark (grant to ADB), and iPSYCH.

## Author contributions

- Analysis: FKS (primary), RKW, TS, EMW, FL, DD, JAK, JG, DSP, JBM
- Sample processing: FKS, RKW, CS, JB-G, MB-H, MN, OM, DMH, TMW, PBM, ADB, iPSYCH-Broad Consortium
- Core PI group: MN, OM, EBR, DMH, TMW, PBM, BMN, ADB, MJD
- Project direction: BMN, ADB, MJD (primary)
- Core writing group: FKS, MJD

## Supplementary Figures / Tables

See accompanying file

## Supplementary Data

Supplementary Data will be available shortly from http://ipsych.au.dk/downloads/

